# Constitutive IFNα protein production in bats

**DOI:** 10.1101/2021.06.21.449208

**Authors:** Vincent Bondet, Maxime Le Baut, Sophie Le Poder, Alexis Lécu, Thierry Petit, Rudy Wedlarski, Darragh Duffy, Delphine Le Roux

**Affiliations:** Translational Immunology Lab, Institut Pasteur, Paris, France; BioPôle Alfort, Ecole Nationale Vétérinaire d’Alfort, Maisons-Alfort, F-94700, France; Anses, INRAE, Ecole Nationale Vétérinaire d’Alfort, UMR VIROLOGIE, Laboratoire de Santé Animale, Maisons-Alfort, F-94700, France; Parc Zoologique de Paris, Muséum National d’Histoire Naturelle, 53 Avenue de Saint-Maurice, F-75012, Paris, France; Parc Zoologique de La Palmyre, 6 Avenue de Royan, F-17570 Les Mathes, France; Bioparc Zoo de Doué La Fontaine, 103 Rue de Cholet, F-49700 Doué-la-Fontaine, France; Anses, INRAE, Ecole Nationale Vétérinaire d’Alfort, UMR BIPAR, Laboratoire de Santé Animale, Maisons-Alfort, F-94700, France

**Author notes:** These authors contributed equally to this work (Vincent Bondet, Maxime Le Baut). These authors contributed equally to this work (Darragh Duffy, Delphine Le Roux). Corresponding authors*: Darragh Duffy, Translational Immunology Lab, Institut Pasteur, 25, rue du Dr. Roux, 75724 Paris Cedex 15, France. Delphine Le Roux, UMR BIPAR, BioPôle Alfort, Ecole Nationale Vétérinaire d’Alfort, 7 avenue du Général de Gaulle, 94700 Maisons-Alfort, France.

**Keywords:** Chiroptera, bats, type I IFN, Simoa digital ELISA, protein levels, antiviral immunity

## Abstract

Bats are the only mammals with self-powered flight and account for 20% of all extant mammalian diversity. In addition, they harbor many emerging and reemerging viruses, including multiple coronaviruses, several of which are highly pathogenic in other mammals, but cause no disease in bats. How this relationship between bats and viruses exists is not yet fully understood. Existing evidence supports a specific role for the innate immune system, in particular type I interferon (IFN) responses, a major component of antiviral immunity. Previous studies in bats have shown that components of the IFN pathway are constitutively activated at the transcriptional level. In this study, we tested the hypothesis that the type I IFN response in bats is also constitutively activated at the protein level. For this we utilized highly sensitive Single Molecule (Simoa) digital ELISA assays, previously developed for humans that we adapted to bat samples. We prospectively sampled four non-native chiroptera species from French zoos. We identified a constitutive expression of IFNα protein in the circulation of healthy bats, and concentrations that are physiologically active in humans. Expression levels differed according to the species examined, but was not associated with age, sex, or health status suggesting constitutive IFNα protein expression independent of disease. These results confirm a unique IFN response in bat species that may explain their ability to coexist with multiple viruses in the absence of pathology. These results may help to manage potential zoonotic viral reservoirs and potentially identify new anti-viral strategies.

## 1. Introduction

With more than 1,200 species, representing about 20% of the total diversity of Mammals, bats are among the most abundant, diverse and geographically dispersed vertebrates on the planet. They are the only mammals capable of active flight and present numerous anatomical variations (1–4). Bats also act as reservoirs for a multitude of viruses, some recognized as highly pathogenic to humans and animals (5–10). Moreover, bats have been shown to be involved in the emergence and reemergence of numerous highly pathogenic zoonotic viruses such as Rhabdoviridae, Paramyxoviridae (Nipah and Hendra viruses), Filoviridae (Ebola and Marburg viruses), and Coronaviridae (11–19) for which they are suspected to be involved in the emergence of the original SARS-CoV-2 viral strain (20). Moreover, most of the bats experimentally infected with viral doses of Hepinaviruses or Lyssaviruses, which are lethal to other mammals, did not show apparent clinical signs. (21–23). It is likely that viruses and their bat hosts have undergone a long process of co-evolution that began several million years ago with the appearance of the first Chiroptera (9, 24, 25). These mechanisms, along with their specific characteristics including longevity, migratory activity, active flight, and population density, may have shaped both the bat immune system and their ability to thwart host responses to viruses, resulting in a balance between persistent infection and absence of pathophysiology (5, 6, 10, 26, 27). One hypothesis is that bats are able to control viral replication through the existence of specific innate antiviral mechanisms (28). Among them, the production of IFN is known to be the first line of defence against viral infections (29) and there is evidence of a strong constitutive genomic expression of type I IFN, mainly of IFNα, in at least two species of Chiroptera *(P. alecto* and *C. brachyotis)* (30). This difference is unique since it has not been observed in other mammals, however, it remains to be confirmed whether this constitutive expression occurs at the protein level.

We have previously used ultra-sensitive Single Molecule Array (Simoa) digital ELISA to measure IFNα protein in the serum of human patients with autoimmune diseases or viral infection whose levels were previously undetectable with conventional ELISA techniques (31). This ultra-sensitive technique is therefore capable of measuring cytokines at very low concentrations in biological fluids, which were previously only measured indirectly by detection of downstream gene induction. Thus, the measurement of IFNα protein in Chiroptera may confirm the hypothesis raised by Zhou *et al.* which is based solely on mRNA measurements and not on direct protein quantification (30). Indeed, constitutively expressed type I IFN mRNA in bats could represent a “ready to use” pool during viral infection, or it could be directly translated into protein resulting in high blood concentrations and viral protection. In this study, we quantified bat IFNα2 protein using ultrasensitive digital ELISA on plasma from four species of captive bats: *P. rodricensis, P lylei, R. aegyptiacus* and *E. helvum.* We also correlated the IFNα protein expression to the corresponding mRNA levels. This study provides new evidence of how the unique immune system of bats may control viruses in the absence of disease and in doing so constitute a constant viral reservoir for zoonotic transmission and potential new pandemics.

## 2. Materials and Methods

### 2.1 Bat cohort and sampling

Four bat species from four French zoos were sampled, during their annual sanitary examination, by the resident veterinary doctor. 0.2 to 0.5 mL of blood was drawn from different veins depending on the species and collected in an EDTA containing tube using a 1mL syringe and a 25G x 5/8 needle (all from Beckton Dickinson, France). Under general anesthesia (O_2_ at 1.5L/min and 5% isoflurane for induction and 2% to maintain the anesthesia) blood of *P. rodricensis* and *R. aegyptiacus* was taken from the medial vein and jugular vein, respectively. *E. helvum* and *P. lylei* were vigilant during blood sampling, which was done from the medial vein for both species. The demographic characteristics and the origins of this cohort are indicated in Table 1. Blood was then split into PAXGene tubes (PreAnalytix GmbH, Qiagen, France) for RNA extraction and RT-qPCR analysis, and Eppendorf tubes to obtain plasma. PAXGene tubes were stored at −20°C until extraction, while Eppendorf tubes were centrifuged at 2500rpm for 10min. Plasma was then removed and stored at −80°C until Simoa analysis.

**Table 1.** Bat species sampled from 4 different zoos.

| Number of individuals | Sex, (female) | Species | Origin |
| --- | --- | --- | --- |
| 108 | 36 (43%) | *Pteropus rodricensis*: 21 (19%) | Parc Zoologique de La Palmyre |
|  |  | *Rousettus aegyptiacus*: 10 (9.3%) | Parc Zoologique de La Palmyre |
|  |  | *Eidolon helvum*: 23 (21%) | Parc Zoologique de Paris |
|  |  | *Pteropus lylei*: 54 (50%) | Bioparc Zoo de Doué La Fontaine |

### 2.2 Bat cell stimulation assays

Bat epithelial cells from Tb 1 Lu cell line (ATCC, United States) were cultured at 37°C, 5% CO_2_, in complete culture medium composed of MEM Eagle medium with 2 mM Glutamine supplemented with 50 U/mL penicillin, 50 μg/mL streptomycin (all from Lonza, Belgium) and with 10% of decomplemented fetal calf serum (Gibco, Thermo Fisher Scientific, France). Before stimulation, cells were plated in 1mL of complete medium per well of 24 well plates and maintained at 37°C, 5% CO_2_ until they reached 2×10^6^ cells/well. Supernatants were removed and 1mL of complete medium including 500HAU/mL mouse influenza virus (Strain H1N1 A/PR81934) was added to the cells or not (unstimulated control). Before stimulation and 1 hour, 3.5 hours, and 23 hours after stimulation, supernatants were sampled and frozen at −80°C for IFNα protein quantification.

### 2.3 Production of the *Rousettus aegyptiacus* IFNα protein for ELISA calibration

The *Rousettus aegyptiacus* IFNα DNA sequence was obtained from a previously published study (32). Nucleotide bases that correspond to the signal peptide were removed, a start codon, spacers, and codons for a 6His tag and a TEV cleavage site were added in the 5’ termination. The cDNA coding for the recombinant protein was chemically-synthesized with optimization for expression in *Escherichia coli.* The recombinant gene was then introduced in a pT7 expression plasmid under the control of a Lac operator and harboring kanamycin resistance. *E. coli* strains were transformed and kanamycin-resistant clones were selected. After optimization, protein production was done, culturing the selected clones in a Luria-Bertani kanamycin (LBkan) medium at 16°C during 16 hours after induction with 1mM isopropyl β-D-1-thiogalactopyranoside (IPTG). Bacteria were then harvested by centrifugation. The pellet was lysed and the soluble extract was obtained after a second centrifugation. This soluble extract was directly used at different dilutions as a calibrator for the digital ELISA assay. It was aliquoted and stored at −80°C before use.

### 2.4 Sample preparation for IFNα2 digital ELISA assay

All plasma samples were first thawed and centrifuged at 10.000g, +4°C for 10 minutes to remove debris. Because bats can harbor many viruses, supernatants were treated in a P2 laboratory for viral inactivation using a standard solvent/detergent protocol used for human blood plasma products (33, 34) and described in (35) and in (36). Briefly, samples were treated with Tri-*n*-Butyl Phosphate (T*n*BP) 0.3% (v/v) and Triton X100 (TX100) 1% (v/v) for 2 hours at room temperature. After treatment, T*n*BP was removed by passing the samples through a C18 column (Discovery DSC-18 SPE from Supelco). For digital ELISA assays, inactivated samples and stimulated cell supernatants were diluted in the Detector / Sample Diluent (Quanterix) added with NP40 0,5% (v/v). They were then incubated for one hour at room temperature before analysis. Global dilution factor was generally 1/6 for plasma samples and 1/3 for stimulated cell supernatants depending on the amount of material available and to allow the optimal protein detection.

### 2.5 IFN«2 digital ELISA assay

The Simoa IFNα2 assay was developed using the Quanterix Homebrew kit and described in (36). The BMS216C (eBioscience) antibody clone was used as a capture antibody after coating on paramagnetic beads (0.3mg/mL), and the BMS216BK biotinylated antibody clone was used as the detector at a concentration of 0.3ug/mL. The SBG revelation enzyme concentration was 150pM. The assay follows a 2-step ELISA configuration. Two calibrators were used; recombinant human IFNα2c (hIFNα2c) purchased from eBioscience and *Rousettus aegyptiacus* IFNα (bIFNα) produced in *Escherichia coli* for this study. The limit of detection (LOD) was calculated by the mean value of all blank runs + 2SD after log conversion.

### 2.6 RNA extraction and IFNα RT-qPCR

Whole blood RNA was extracted manually from PAXGene tubes, following manufacturer’s instructions (Blood RNA extraction kit, Qiagen, France). After extraction, samples were inactivated at 65°C for 5min then stored at −80°C until RT-qPCR. RT-qPCR was done using the qScript XLT One-Step RT-qPCR mix following manufacturer’s instructions (Quanta BioSciences, Inc., United States). Taqman probes (Applied Biosystems, ThermoFisher Scientific, France) and primers (Eurofins, France) for IFN-α1, IFNα2 and IFNα3 were described previously for *P. alecto* bat species in (30) and used here. Probes and primers for the bat GAPDH housekeeping gene were designed using Primer-BLAST from NCBI (https://www.ncbi.nlm.nih.gov/tools/primer-blast/) and are presented in table S2. All data were normalized relative to the housekeeping gene (GAPDH) as indicated. The expression level of the target genes was calculated using the standard curve method and expressed as copy numbers relative to the housekeeping gene.

### 2.7 Nested RT-qPCR for pan-coronaviruses in mRNA from bat whole blood

RT-qPCR for potential coronaviruses in bat whole blood RNA was performed from the mRNA extracted previously and following the protocol previously published in (37).

### 2.8 Statistical analyses

GraphPad Prism 8 was used for statistical analysis. Mann-Whitney tests were used to compare two groups such as female and male, or healthy and disease. ANOVA tests (Kruskal-Wallis) with Dunn’s post testing for multiple comparisons were used to test for differences between multiple bat species. For all analyses, p values less than 0.05 were considered statistically significant, with *p< 0.05; **p<0.01; ***p<0.001; ****p<0.0001. Median values were reported on figures. Spearman correlations are used to compare continuous variables such as mRNA level or age and protein production.

### 2.9 Data availability

All available data from the bat cohort are shown in Supplementary Table S1.

## 3. Results

### 3.1 Bat IFNα protein detection with an IFNα2 digital ELISA assay

Anti-bat IFNα antibodies are not commercially available for the development of bat-specific IFNα ELISA. However, given the ultra-sensitivity of human IFNα digital ELISA which detects protein at attomolar concentrations (31), and potential cross-species reactivity, we hypothesized that our existing human assay could also detect bat IFNα. As a first proof of concept, we stimulated a bat lung epithelial cell line with influenza virus (Strain H1NI A/PR81934) and tested the recovered supernatant with a human IFNα2 digital ELISA. We observed a significant induction of IFNα2 protein at 1hr and 3.5 hrs as compared to the unstimulated control (Fig 1a).

**Figure 1.**
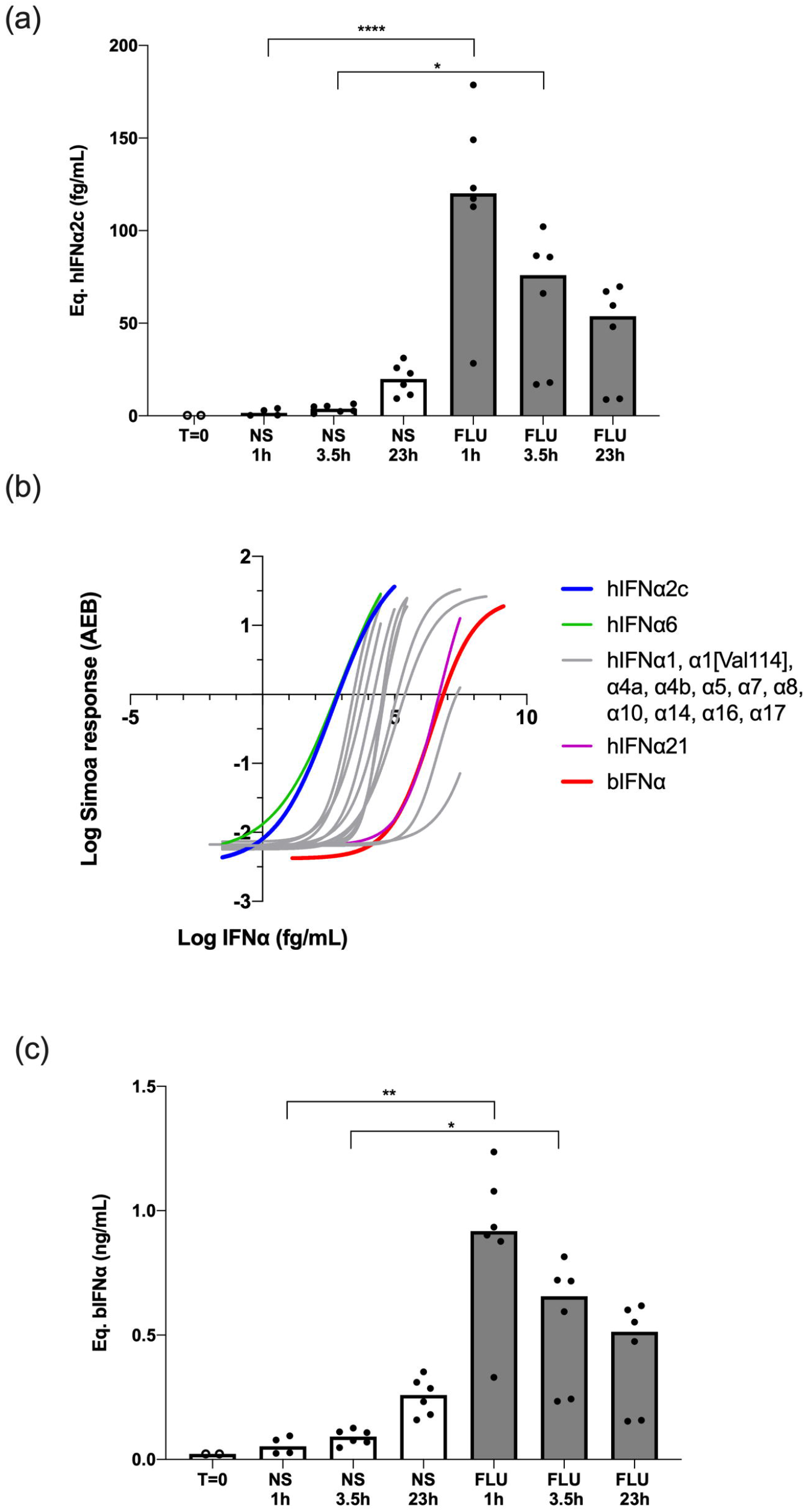
Bat IFNα proteins are detected with a human IFNα2 digital ELISA assay. **(a)** IFNα protein levels expressed as equivalent human IFNα2c concentrations obtained after stimulation of bat epithelial cells Tb 1 Lu with 500HAU/mL mouse influenza virus (FLU) or unstimulated (NS) for 0 to 24 hours at 37°C and 5% CO_2_. **(b)** IFNα2 digital ELISA assay response (AEB) as a function of the IFNα concentration for the *Rousettus aegyptiacus* IFNα calibrator (bIFNα) produced in *Escherichia coli* (red) in comparison with the human IFNα2c calibrator (blue) and the 12 other human IFNα subtypes. **(c)** IFNα protein levels expressed as equivalent bIFNα concentrations obtained after stimulation of bat epithelial cells as described previously. Box plots represent median and individual values represented by dots are reported on figures, a & c representing pooled results from 3 independent experiments

These initial results were extrapolated from a standard curve of a human recombinant IFNα2 protein. To better adapt our assay to bat species, we produced a recombinant *Rousettus aegyptiacus* IFNα protein (bIFNα) from *Escherichia coli* competent bacteria. SDS-Page analysis of the soluble and insoluble extracts obtained from the bacteria pellet showed that bIFNα was mainly produced as an insoluble form even at low induction temperature (Fig S1a). Comparing profiles before and after induction of the protein expression, SDS-Page analysis showed that the unique bIFNα band appeared alone at this mass (Fig S1a). Western-Blot analysis of the soluble fraction after induction at 16°C using the IFNα2 assay detection antibody revealed that the protein was expressed in a single band at the expected molecular weight (Fig S1a). The purification from the soluble extract failed: the bIFNα protein was not selected at the expected molecular weight (Fig S1b) and the western-Blot analysis revealed no affinity at the purified molecular weight (Fig S1a). The purification from the insoluble extract succeeded, but the renaturation of the protein failed (Fig S1c). So we used the soluble extract itself as a calibrator after quantification of bIFNα. Global protein quantification of the soluble extract was done using the BCA assay, and the bIFNα protein concentration in the soluble extract was estimated after gel densitometry for potential use as a digital ELISA calibrator.

To explore the ability of the IFNα2 digital ELISA assay designed for the quantification of human interferons, to quantify bat IFNα species, we compared the responses of the assay to bIFNα protein and all 13 human IFNα subtypes (Fig. 1b). As expected the assay revealed a weaker response for bIFNα in comparison with hIFNα2c. However, the affinity of the human mAb for the bat protein was comparable to the human subtypes, with bIFNα and hIFNα21 showing very similar affinities, and two human species showing weaker responses (Fig. 1b). Using the bIFNα protein as the calibrator, we re-calculated the cellular response after *in vitro* influenza stimulation and observed similar results with the highest concentrations present after 1hr of influenza stimulation (Fig 1c). The only difference observed was related to the scale of these results, due to the lower affinity of the mAb for the bIFNa2 calibrator.

To better understand the cross-species reactivity we compared the sensitivities of the 13 IFNα subtypes as previously described (36), the 5 other human IFNβ, IFNλ1, IFNλ2, IFNω and IFNγ, and the 5 mouse IFNα1, IFNα3, IFNα4, IFNα11 and IFNα13, with their available online sequences in the UniProtKB database (www.uniprot.org) after alignment using the CLUSTALW software (www.genome.jp/tools-bin/clustalw). We also considered the fact that epitopes must be accessible to the antibodies and so studied the IFNα2, IFNα14 and IFNω three-dimensional structures published online in the PDB database (www.rcsb.org) after spacial alignment using the PyMOL software (www.pymol.org). This analysis suggests that the epitope recognized by the IFNα2 mAb could be the ^110^LMKED sequence in the human IFNα2 molecule. *Rousettus aegyptiacus* IFNα and *Pteropus rodricensis* IFNα1, IFNα2 and IFNα3 protein sequences were previously described by Omatsu *et al* (32)and Zhou *et al.* (30) respectively. Alignment of these molecules showed that the corresponding amino-acids in the IFNα2 assay epitope position are LMNED for the 3 IFNα species from *Pteropus* and LLDED for *Rousettus* (Fig S2a). The LMNED sequence also appears in human IFNα16 and IFNα17, two species for which the IFNα2 assay shows a positive response. The M-→L substitution concerns two apolar amino-acids. The K-→D substitution changes a positive with a negative charged amino-acid, but these two amino-acids are then hydrophilic and so do not produce a detrimental α-helix coil in the structure. This *in silico* analysis provides support for how the IFNα2 antibody assay may recognize *Rousettus* and *Pteropus* IFNα protein.

### 3.2 IFNα proteins are constitutively elevated in plasma of bat species

Having validated the assay for its ability to detect bat IFNα, we analyzed plasma samples from 4 bat species sampled from French zoos (Table 1) with the digital ELISA assay. Results are presented using the two calibrators; hIFNα2c (Fig. 2a) and bIFNα (Fig. 2b). A greater number of samples were above the assay limit of detection using the bat protein calibrator as compared to the human protein, confirming the interest of using a bat specific protein. IFNα protein responses obtained within species were relatively consistent. *Pteropus rodricensis* and *Rousettus aegyptiacus* showed significantly elevated IFNα protein plasma levels as compared to *Eidolon helvum* and *Pteropus lylei,* with both the hIFNα2c (p<0.05) and bIFNα (p<0.0001) calibrators. Notably in *Eidolon helvum* plasma IFNα was largely undetectable with both assays revealing interesting inter-species differences. When available, we assessed whether age, sex, or presence of mild clinical symptoms (Table S1) were associated with IFNα protein levels in all species but found no significant associations (Fig S3). We also tested for presence of corona viruses in the blood but found no evidence (data not shown). These results support the hypothesis that certain bat species have physiological levels of circulating IFNα protein in healthy conditions.

**Figure 2.**
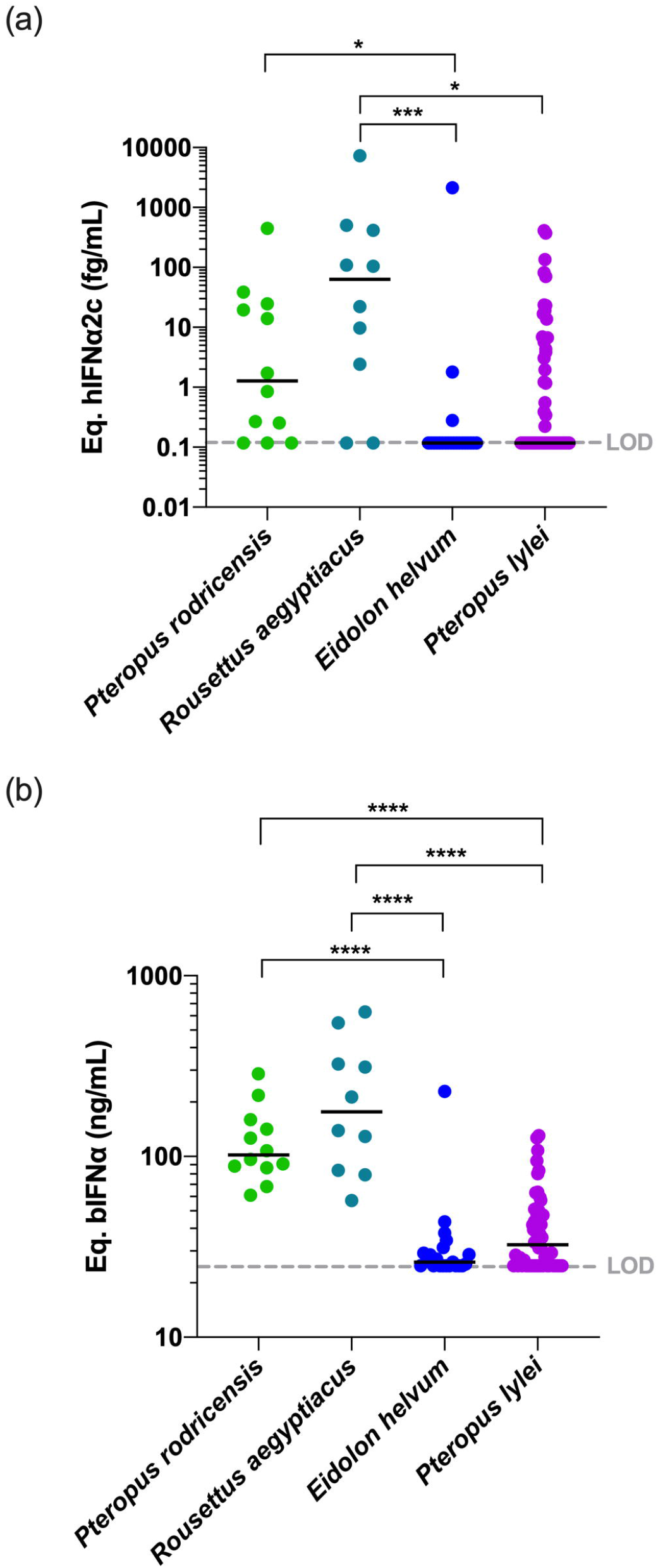
IFNα protein concentrations in plasma of four bat species. IFNα concentrations measured in plasma from four bat species expressed **(a)** as hIFNα2c or **(b)** as bIFNα equivalent concentrations. LOD is the limit of detection level of the assay. Median represented by black line with individual animals shown by colour coded dots. Kruskal-Wallis test with Dunn’s post testing for multiple comparisons was used, *p<0.05, ***p<0.0001, ****p<0.001. *P rodricensis* (green, n=12) *R. aegyptiacus* (light blue, n=10), *E. helvum* (dark blue, n=21) and *P. lylei (purple, n=52).*

### 3.3 IFNα mRNA are constitutively expressed in bat leukocytes

The constitutive mRNA expression of bat IFNα genes has been previously described for *Rousettus aegyptiacus* (32) and *Pteropus rodricensis* (30). To test whether the protein plasma levels we observed were linked with leukocyte mRNA expression, and to extend these observations to *Eidolon helvum* and *Pteropus lylei,* we quantified gene expression of IFNα1, IFNα2 and IFNα3 in our cohort using RT-qPCR and normalizing the results using GAPDH mRNA. Results from *Rousettus aegyptiacus* were negative, perhaps explained by a lack of specificity of the primers utilized. In the other bat species examined the number of IFNα mRNA copies were globally similar to GAPDH, supporting that IFNα species are constitutively expressed at the mRNA level. The IFNα2 subtype mRNA was the most expressed, while IFNα3 was the most variable between species. *Eidolon helvum* had the lowest IFNα mRNA levels, reflecting the absence of detectable protein in this species. *Pteropus lylei* had overall the highest IFNα RNA levels. Significant differences (p<0.05) were observed between *Eidolon helvum* and *Pteropus lylei* for IFNα2 and IFNα3, and between *Pteropus rodricensis* and *Pteropus lylei* for IFNα3. This also indicates that high levels of variation could be observed within a same genus, and also within the same species.

### 3.4 IFNα mRNA expression and protein production could be linked within each species

While *Eidolon helvum* showed lower mRNA and protein levels as compared to the other species, and *Pteropus rodricensis* medium mRNA levels and a high protein level, *Pteropus lylei* showed the highest RNA and the lowest protein levels (Fig. 2a, b and Fig. 3a). Therefore, we observed no overall direct link between IFNα mRNA expression and protein plasma concentrations across all species examined. To directly explore this hypothesis, we tested the correlations for each gene and protein for *Pteropus rodricensis* samples where sufficient detectable measurements were available for both parameters. In this relatively small sample size (n=8 for paired samples) we observed a strong statistically significant correlation (Rs=0.952, p=0.001) between IFNα2 RNA and protein levels (Fig. 3b). No correlation was observed with IFNα1 and IFNα3 genes (Fig. 3b). This result may reflect that the ELISA assay is designed for the IFNα2 subtype.

**Figure 3.**
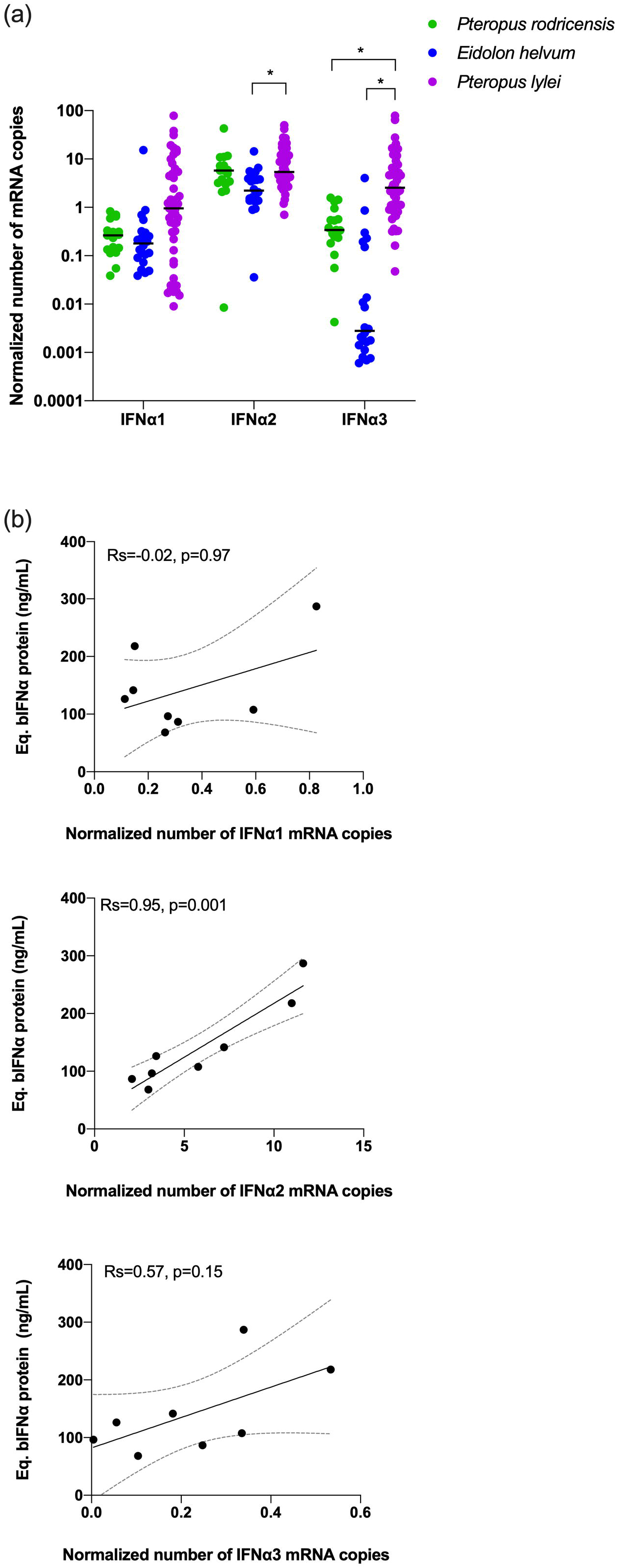
IFNα mRNA levels in bats and mRNA-protein correlations. **(a)** Number of copies measured in whole blood for IFNα1, IFNα2 and IFNα3 mRNAs as normalized to GAPDH in three bat species. Probes used here were not suitable for *Rousettus aegyptiacus*. Median represented by black line with individual animals shown by colour coded dots. Kruskal-Wallis test with Dunn’s post testing for multiple comparisons was used. *p<0.05. **(b)** Correlation plots between the Eq. bIFNα protein concentration obtained using the IFNα2 digital ELISA assay and the GAPDH-normalized number of IFNα1, IFNα2 and IFNα3 mRNA copies. Spearman method is used for correlation analysis with Spearman’s Rank Correlation Coefficient R (Rs) and p values reported (n=8).

## 4. Discussion

Type I interferons trigger the downstream activation of hundreds of critical genes as part of the antiviral immune response. Because of this potency, IFN proteins are secreted at relatively low concentrations as compared to other major cytokines, and multiple regulatory mechanisms exist to control their effects (38). Potential negative consequences of over activation of interferon responses is reflected by their direct implication in multiple autoimmune conditions such as lupus and interferonopathies (39). Furthermore, long-term treatment of patients (ie chronic HCV patients) with type I IFN based therapies can induce serious side effects including depression (40).

Because of these potential negative effects of IFN signaling, recent studies that reported constitutive IFN gene expression in healthy bats were unexpected. However, it provided strong evidence for how bats may potentially live healthily with multiple viral species that are pathological to other mammals including humans. Nevertheless, it raised many additional questions, namely the one which we addressed in our study as to whether IFN protein is also constitutively elevated in bats. This question is not so trivial to address for multiple reasons. IFN proteins have been challenging to directly quantify due to their low physiological concentrations in biological samples, and most studies have utilized proxy readouts such as interferon stimulated gene (ISG) expression or cytopathic assays. The development of digital ELISA such as Simoa overcame these challenges as we demonstrated by the measurement of all 13 IFNα human subtypes, and more recently specifically IFNα2 in multiple human cohorts (31, 36, 41). However additional challenges exist for the study of interferon in non-human species, in particular bats, most notably the lack of specific reagents in particular monoclonal antibodies which are required for ELISA technologies. We tested the hypothesis that attomolar digital ELISA sensitivity combined with species cross-reactivity would enable the quantification of bat IFNα protein. After assay validation on virus stimulated bat cell cultures, we further modified the assay with a recombinant bat protein as the standard calibrator. Applying this assay to 4 species of bats conclusively showed constitutive expression of IFNα protein in the circulation of healthy animals. While we cannot yet make a direct comparison, these levels are significantly higher as compared to healthy humans, where IFNα is essentially undetectable (36) even at attomolar sensitivity.

Our study contains some inherent weaknesses. IFNα protein concentrations were calculated using human IFNα2c or *Rousettus aegyptiacus* IFNα protein (bIFNα) produced in *E. coli* as calibrators, with results respectively in the fg/mL and the ng/mL ranges. Such a difference could be due to different antibody specificities, potentially due to incorrect folding of the bIFNα protein produced in *E. coli* strains. This is supported by the observation that the bIFNα protein is mainly expressed in the insoluble fraction even at 16°C (Fig. S1c), that the protein failed to renature after urea purification (Fig. S1b), and that the viral inactivation solvent/detergent protocol had a greater effect on the IFNα2 assay response to bIFNα (>1Log) than hIFNα2c (Fig. S4), suggesting greater insolubility of the bat protein. Additional improvements of the assay could be envisioned such as production of a purer and better folded recombinant protein in mammalian cells, and eventually the production of a bat specific monoclonal antibody against this protein.

Despite these technical limitations, we were able to show elevated levels of plasma IFNα protein in certain bat species, which also correlated with expression levels of the IFNα2 gene. While additional confirmatory experiments will be required, the inter-species differences in plasma IFNα protein is an interesting observation. It would also be interesting in future studies to assess whether these IFNα protein differences have an impact on viral levels and diversity within the different bat species. Lastly, our results raise additional new questions on the nature of bat physiology, in particular how the constitutively activated type I IFN response is maintained in bats without resulting in pathological conditions such as those observed in human autoimmune disease. Chronic IFN activation, in particular during growing and development phases, can have significant neurological effects as observed in interferonopathies such as STING mutation patients (42). Finally given the important role of type I interferon for protection to infection with SARS-CoV-2 (35, 43), and the potential role bats have played in seeding the COVID-19 pandemic (20) understanding this host-virus relationship could have major implications for pandemic preparedness. In summary improved knowledge on the special nature of bat IFN regulation could have major implications for our basic understanding of IFN biology, its continued use as a therapeutic, and our capacity to prepare for viral pandemics.

## Abbreviations

AEB: average enzyme per bead
HAU: hemagglutinating units
IFN: interferon
LOD: limit of detection

## Acknowledgements

We thank the ProteoGenix company (Schiltigheim, France) for the production of the bIFNα ELISA assay calibrator; Marielle Cochet for technical help; Florence Va, Dr Vitomir Djokic and Isabelle Badreau for helping in RT-qPCR datas analysis; the technical staff of the zoos and the Association Française des Vétérinaires de Parcs Zoologiques for their help in bat handling and sampling. We thank the CBUTechS platform of the Institut Pasteur for access to the Simoa instrument. DD acknowledges support from the ANR (grant number CE17001002).

---

**Figure S1. Production and purification of the bIFNα calibrator.**

**(a)** SDS-Page profile obtained before IPTG induction (Ø) and after induction at 16, 30 or 37°C in the soluble or insoluble extract of *Escherichia coli* cells transformed to produce bIFNα. Molecular weights (MW) are indicated in kDa (left panel). Western-Blot profile obtained from the soluble fraction after induction at 16°C (right panel). After SDS-Page migration, proteins were transferred on a nitrocellulose membrane then incubated with the IFNα2 assay detection antibody. Streptavidin-conjugated horseradish peroxydase was added to the membrane and signal was revealed using chemiluminescence. **(b)** SDS-Page profile for each bIFNα purification step from the soluble fraction (left panel) after IPTG induction at 16°C. Column load (IN), column flow-through (FT), washes (W1, W2 and W3) and elution fractions (E1 to E9). The middle panel shows the SDS-Page profile for the pools of eluted fractions. The red arrow indicates the bIFNα protein at its right molecular weight. **(c)** SDS-Page profile for each bIFNα purification step from the insoluble fraction (left panel) after IPTG induction at 16°C as previously described. Right panel shows SDS-Page profile for each renaturation step of the pool of eluted fractions obtained from the insoluble *E. coli* extract: 4M, then 2M, then 1M urea and finally without urea (ØM), the ultimate step where the protein is lost. Red arrows indicate the bIFNα protein at his right molecular weight.

**Figure S2. Bat IFNα proteins have a similar IFNα2 assay epitope to human IFNα proteins**

Protein sequence alignment obtained for human IFNα1, IFNα2, IFNα16 and IFNα17, for *Pteropus alecto* IFNα1, IFNα2 and IFNα3, for *Rousettus aegyptiacus* IFNα, and for mouse IFNα1, IFNα3, IFNα4, IFNα11 and IFNα13 using the CLUSTALW software from position 90 to 130 referred to hIFNα2. The red rectangle highlights the suspected epitope recognized by the antibodies of the IFNα2 assay.

**Figure S3. Age, sex and clinical symptoms were not associated with IFNα protein levels**

**(a)** Correlation plot between age of all studied bat specimens and IFNα protein levels expressed as bIFNα equivalent concentration. Spearman method is used. **(b)** IFNα protein levels expressed as bIFNα equivalent concentration for female and male groups, all species combined. Median represented by black line. Mann-Whitney test is used, ns: p>0.05. **(c)** IFNα protein levels expressed as bIFNα equivalent concentration for healthy and mild clinical symptom groups, all species combined. Median represented by black line. Mann-Whitney test is used, ns: p>0.05.

**Figure S4. Viral inactivation decreases the affinity of the assay for bIFNα but not hIFNα2c.**

IFNα2 assay response (AEB) as a function of IFNα concentrations for hIFNα2c and bIFNα untreated or after viral inactivation.

## Tables Legends

**Table 1. Demographic characteristics and origins of the bat cohort**

Number of individuals, gender, species and origins for the bat cohort. Data are shown as the n (%).

**Table S1. All data available for each sample**

Species, gender, GAPDH-normalized number of IFNα1, IFNα2 and IFNα3 mRNA copies, interferon α concentrations expressed as hIFNα2c or bIFNα equivalent concentrations and notes available for each sample.

**Table S2. Primers and probes sequences**

List of primers and probes used in RT-qPCR for bat IFNα1, IFNα2, IFNα3 and GADPH, based on published data from Zhou et al. 2016 for the bat IFN genes described for *P. alecto* and designed for this study for GADPH, from PrimerBlast and available datas for GADPH gene in *P. alecto* and *R. aegyptiacus* species.

## Notes

### Competing Interest Statement

The authors have declared no competing interest.

## References

1. Teeling EC, et al. (2005) A molecular phylogeny for bats illuminates biogeography and the fossil record. Science 307(5709):580–584.

2. Amador LI, Moyers Arévalo RL, Almeida FC, Catalano SA, & Giannini NP (2018) Bat Systematics in the Light of Unconstrained Analyses of a Comprehensive Molecular Supermatrix. Journal of Mammalian Evolution 25(1):37–70.

3. Simmons NB (2005) Evolution. An Eocene big bang for bats. Science 307(5709):527–528.

4. Sadier A, et al. (2021) Making a bat: The developmental basis of bat evolution. Genetics and molecular biology 43(1 Suppl 2):e20190146.

5. Wang LF, Walker PJ, & Poon LL (2011) Mass extinctions, biodiversity and mitochondrial function: are bats ‘special’ as reservoirs for emerging viruses? Current opinion in virology 1(6):649–657.

6. Calisher CH, Childs JE, Field HE, Holmes KV, & Schountz T (2006) Bats: important reservoir hosts of emerging viruses. Clinical microbiology reviews 19(3):531–545.

7. Wong S, Lau S, Woo P, & Yuen KY (2007) Bats as a continuing source of emerging infections in humans. Reviews in medical virology 17(2):67–91.

8. Luis AD, et al. (2013) A comparison of bats and rodents as reservoirs of zoonotic viruses: are bats special? Proceedings. Biological sciences 280(1756):20122753.

9. Rodhain F (2015) [Bats and Viruses: complex relationships]. Bulletin de la Societe de pathologie exotique 108(4):272–289.

10. Brook CE & Dobson AP (2015) Bats as ‘special’ reservoirs for emerging zoonotic pathogens. Trends in microbiology 23(3):172–180.

11. Halpin K, Young PL, Field HE, & Mackenzie JS (2000) Isolation of Hendra virus from pteropid bats: a natural reservoir of Hendra virus. The Journal of general virology 81(Pt 8):1927–1932.

12. Chua KB, Wang LF, Lam SK, & Eaton BT (2002) Full length genome sequence of Tioman virus, a novel paramyxovirus in the genus Rubulavirus isolated from fruit bats in Malaysia. Archives of virology 147(7):1323–1348.

13. Leroy EM, et al. (2005) Fruit bats as reservoirs of Ebola virus. Nature 438(7068):575–576.

14. Lau SK, et al. (2005) Severe acute respiratory syndrome coronavirus-like virus in Chinese horseshoe bats. Proceedings of the National Academy of Sciences of the United States of America 102(39):14040–14045.

15. Li W, et al. (2005) Bats are natural reservoirs of SARS-like coronaviruses. Science 310(5748):676–679.

16. Towner JS, et al. (2007) Marburg virus infection detected in a common African bat. PloS one 2(8):e764.

17. Towner JS, et al. (2009) Isolation of genetically diverse Marburg viruses from Egyptian fruit bats. PLoS pathogens 5(7):e1000536.

18. Pourrut X, et al. (2009) Large serological survey showing cocirculation of Ebola and Marburg viruses in Gabonese bat populations, and a high seroprevalence of both viruses in Rousettus aegyptiacus. BMC infectious diseases 9:159.

19. Baker KS, et al. (2013) Novel, potentially zoonotic paramyxoviruses from the African straw-colored fruit bat Eidolon helvum. Journal of virology 87(3):1348–1358.

20. Wacharapluesadee S, et al. (2021) Evidence for SARS-CoV-2 related coronaviruses circulating in bats and pangolins in Southeast Asia. Nature communications 12(1):972.

21. Setien AA, et al. (1998) Experimental rabies infection and oral vaccination in vampire bats (Desmodus rotundus). Vaccine 16(11-12):1122–1126.

22. Williamson MM, Hooper PT, Selleck PW, Westbury HA, & Slocombe RF (2000) Experimental hendra virus infectionin pregnant guinea-pigs and fruit Bats (Pteropus poliocephalus). Journal of comparative pathology 122(2-3):201–207.

23. Middleton DJ, et al. (2007) Experimental Nipah virus infection in pteropid bats (Pteropus poliocephalus). Journal of comparative pathology 136(4):266–272.

24. Badrane H & Tordo N (2001) Host switching in Lyssavirus history from the Chiroptera to the Carnivora orders. Journal of virology 75(17):8096–8104.

25. Drexler JF, et al. (2012) Bats host major mammalian paramyxoviruses. Nature communications 3:796.

26. Swanepoel R, et al. (2007) Studies of reservoir hosts for Marburg virus. Emerging infectious diseases 13(12):1847–1851.

27. Schountz T, Baker ML, Butler J, & Munster V (2017) Immunological Control of Viral Infections in Bats and the Emergence of Viruses Highly Pathogenic to Humans. Frontiers in immunology 8:1098.

28. Baker ML, Schountz T, & Wang LF (2013) Antiviral immune responses of bats: a review. Zoonoses and public health 60(1):104–116.

29. de Weerd NA & Nguyen T (2012) The interferons and their receptors--distribution and regulation. Immunology and cell biology 90(5):483–491.

30. Zhou P, et al. (2016) Contraction of the type I IFN locus and unusual constitutive expression of IFN-alpha in bats. Proceedings of the National Academy of Sciences of the United States of America 113(10):2696–2701.

31. Rodero MP, et al. (2017) Detection of interferon alpha protein reveals differential levels and cellular sources in disease. The Journal of experimental medicine 214(5):1547–1555.

32. Omatsu T, et al. (2008) Induction and sequencing of Rousette bat interferon alpha and beta genes. Veterinary immunology and immunopathology 124(1-2):169–176.

33. Horowitz B, et al. (2004) WHO Expert Committee on Biological Standardization. World Health Organization technical report series 924:1–232, backcover.

34. Kuhnel D, et al. (2017) Inactivation of Zika virus by solvent/detergent treatment of human plasma and other plasma-derived products and pasteurization of human serum albumin. Transfusion 57(3pt2):802–810.

35. Hadjadj J, et al. (2020) Impaired type I interferon activity and inflammatory responses in severe COVID-19 patients. Science 369(6504):718–724.

36. Bondet V, et al. (2021) Differential levels of IFNalpha subtypes in autoimmunity and viral infection. Cytokine:155533.

37. Ar Gouilh M, et al. (2018) SARS-CoV related Betacoronavirus and diverse Alphacoronavirus members found in western old-world. Virology 517:88–97.

38. McNab F, Mayer-Barber K, Sher A, Wack A, & O’Garra A (2015) Type I interferons in infectious disease. Nature reviews. Immunology 15(2):87–103.

39. Picard C & Belot A (2017) Does type-I interferon drive systemic autoimmunity? Autoimmunity reviews 16(9):897–902.

40. Hauser P (2004) Neuropsychiatric side effects of HCV therapy and their treatment: focus on IFN alpha-induced depression. Gastroenterology clinics of North America 33(1 Suppl):S35–50.

41. Rodero MP, et al. (2017) Type I interferon-mediated autoinflammation due to DNase II deficiency. Nature communications 8(1):2176.

42. Fremond ML & Crow YJ (2021) STING-Mediated Lung Inflammation and Beyond. Journal of clinical immunology 41(3):501–514.

43. Bastard P, et al. (2020) Autoantibodies against type I IFNs in patients with life-threatening COVID-19. Science 370(6515).

