## Supplementary figures and images for "Constitutive IFNα protein production in bats"

### Supplemental Figure 1

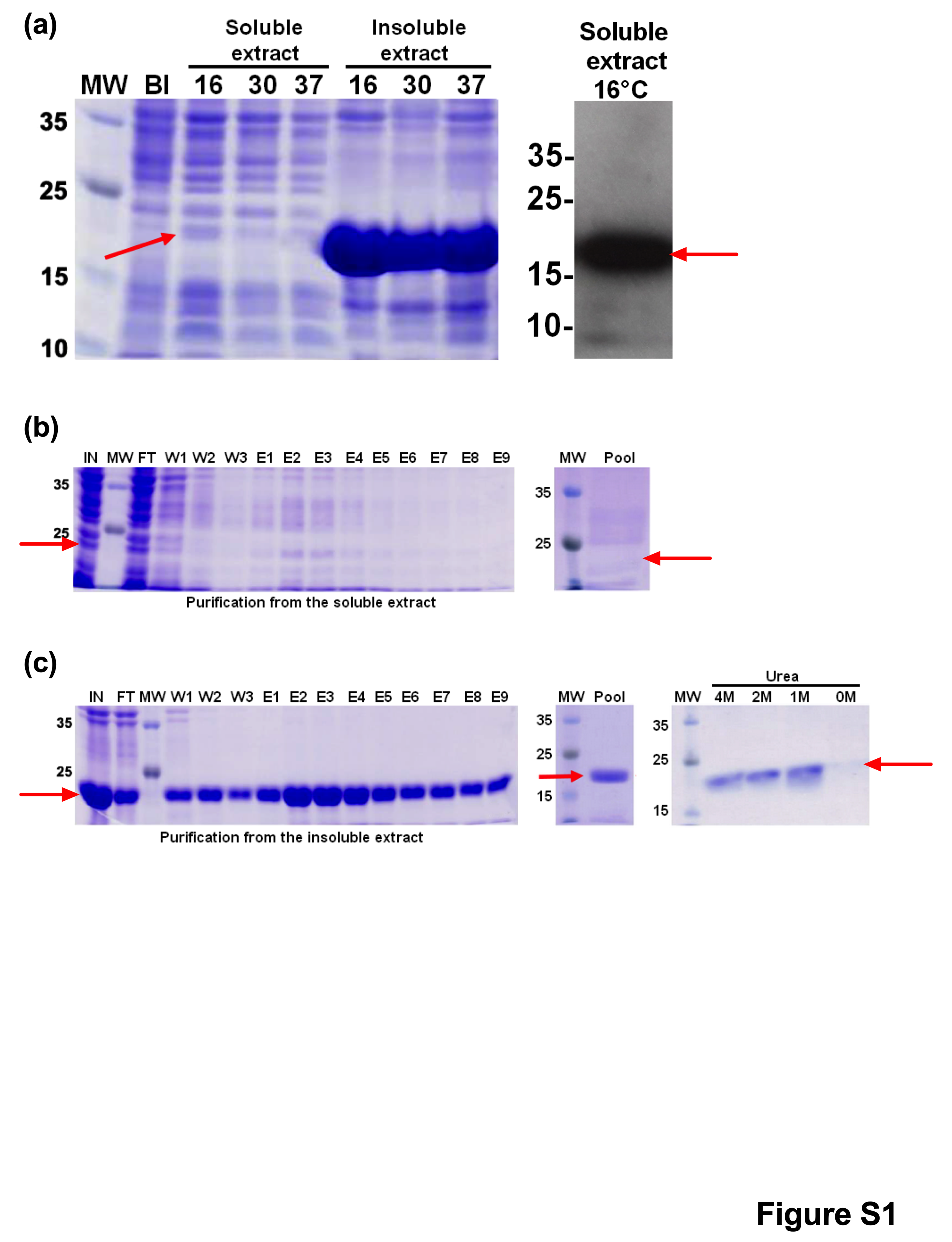

### Supplemental Figure 2

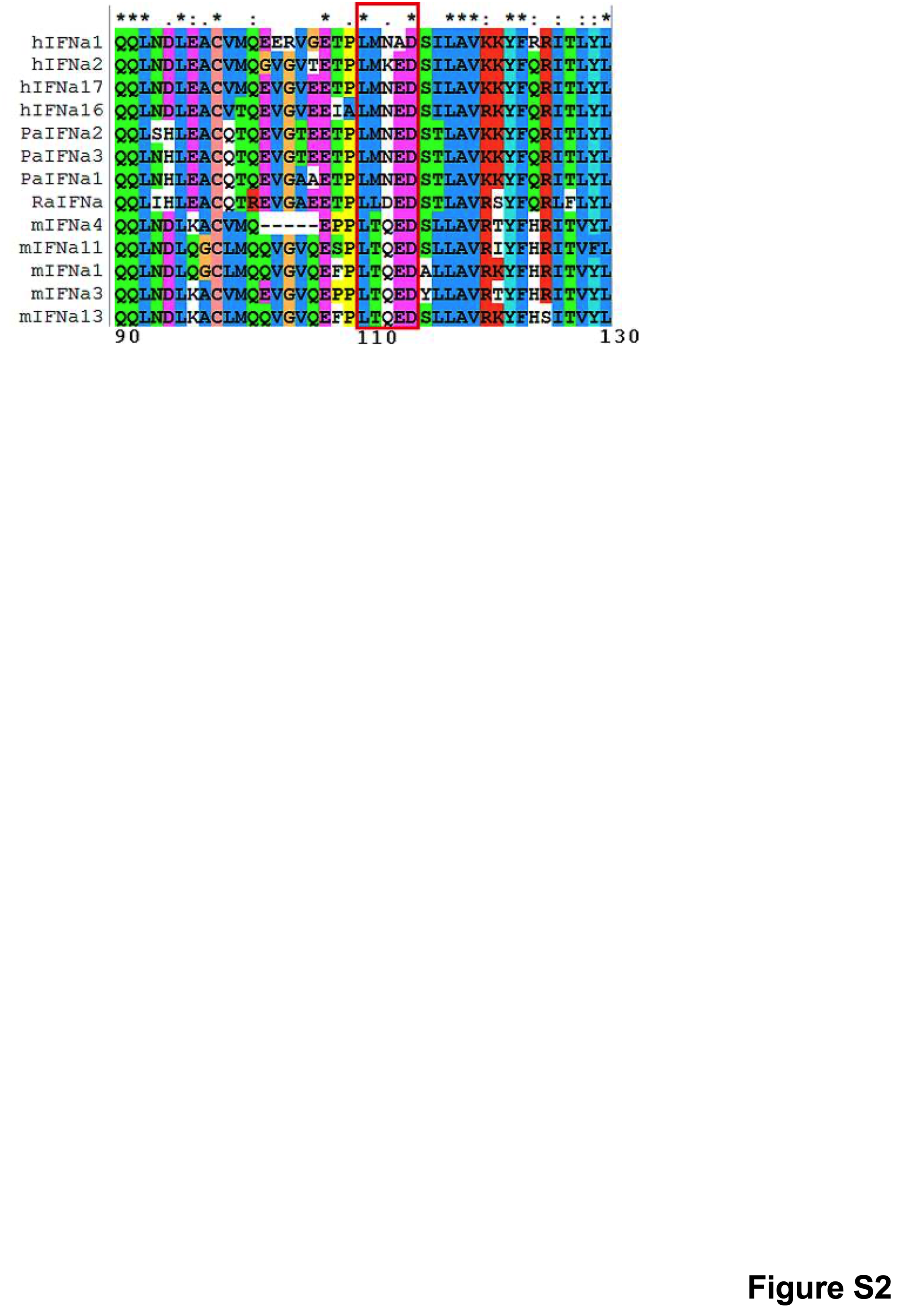

### Supplemental Figure 3

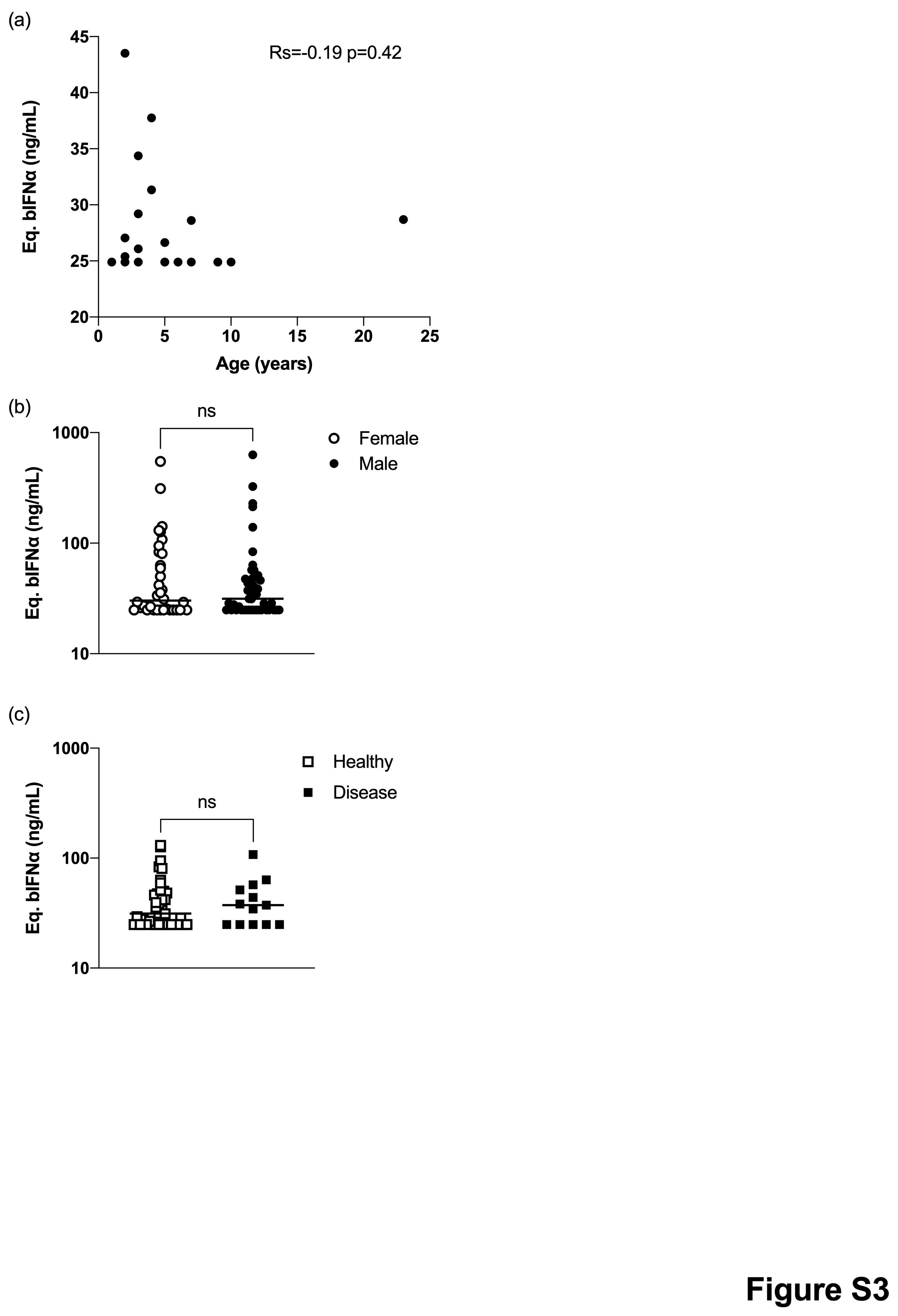

### Supplemental Figure 4

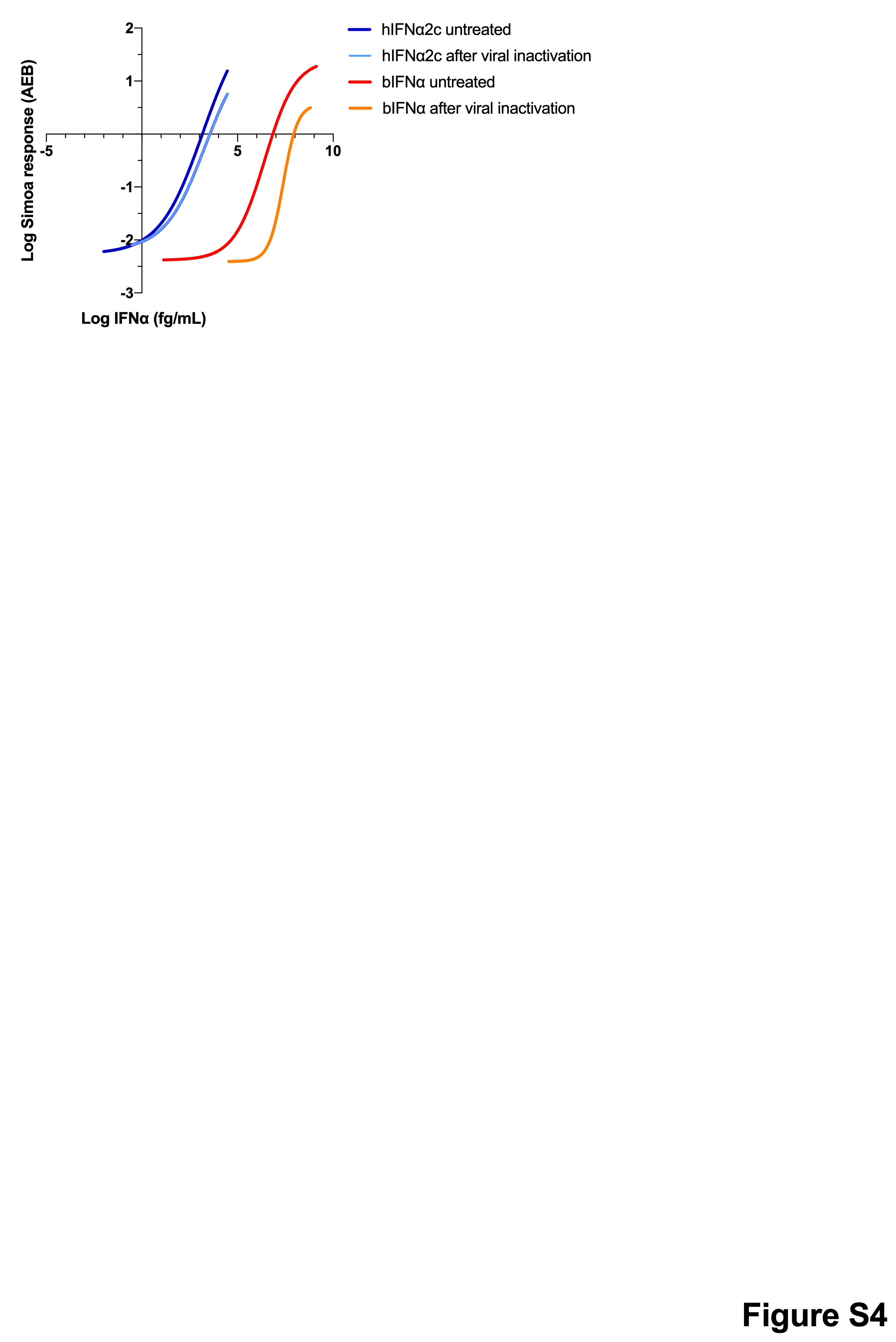
